# Rapid environmental change in games: complications and curiosities

**DOI:** 10.1101/397497

**Authors:** Pete C Trimmer, Brendan J Barrett, Richard McElreath, Andrew Sih

## Abstract

Human-induced rapid environmental change (HIREC) has recently led to alterations in the fitness and behavior of many organisms. Game theory is an important tool of behavioral ecology for analyzing evolutionary situations involving multiple individuals. However, game theory bypasses the details by which behavioral phenotypes are determined, taking the functional perspective straight from expected payoffs to predicted frequencies of behaviors. In contrast with optimization approaches, we identify that to use existing game theoretic models to predict HIREC effects, additional mechanistic details (or assumptions) will often be required. We illustrate this in relation to the hawk-dove game by showing that three different mechanisms, each of which support the same ESS prior to HIREC (fixed polymorphism, probabilistic choice, or cue dependency), can have a substantial effect on behavior (and success) following HIREC. Surprisingly, an increase in the value of resources can lead to a reduction in payoffs (and vice versa), both in the immediate- and long-term following HIREC. An increase in expected costs also increases expected payoffs. Along with these counter-intuitive findings, this work shows that simply understanding the behavioral payoffs of existing games is insufficient to make predictions about the effects of HIREC.

It’s the little details that are vital. Little things make big things happen.

John Wooden

## Introduction

Human-induced rapid environental change (HIREC) poses a threat to the persistance of many species and populations through factors such as climate change, habitat loss or fragmentation, increased human harvesting, exposure to novel biotic (e.g., predators, competitors, pathogens) or abiotic (e.g., pollutants) stressors and/or availability of novel, inappropriate resources. A key aspect of organismal response to HIREC is their initial behavioral response [1,2]. Whilst some animals exhibit adaptive behavioral responses to HIREC [3,2], others show maladaptive behaviors, falling into evolutionary or ecological traps [4,5]. If a population survives the intial selective pressure of a rapid environmental change, then the species can potentially evolve to cope better with HIREC. Thus, initial behavioral responses to changes are critically important for species persistence. Despite much empirical study, few attempts have been made to use explicit mathematical theory to explain the variation in initial response to HIREC (but see [6]). The key challenge is to identify the rationale and then develop models that explain both adaptive and maladaptive responses to HIREC [7].

Based on the evolutionary trap literature, a hypothesis is that how organisms will behave following HIREC depends on cue-response systems that they evolved in their ancestral settings [8]. Models that predict responses to an aspect of HIREC first specify the general mechanism by which cues govern behavior (and typically set parameters of that system as though they have been fine-tuned by natural selection). This approach then posits that immediately after HIREC, organisms use their previously adaptive cue-response systems to respond to the novel situation (e.g., [6]). This approach can then identify how evolutionary match versus mismatch between the past conditions that shaped a cue-response system and the novel conditions after HIREC explains variation in performance after HIREC [7, 8, 6]. A perhaps obvious, implicit assumption is that if the organism’s previously adaptive cue-response system produces adaptive responses to HIREC, it will enjoy higher fitness than if it exhibits a fixed strategy (no adjustment to HIREC) or a less-than optimal plastic response to HIREC. To emphasize, this approach assumes that individuals exhibit behavioral plasticity guided by previously adaptive cue-response systems, but that the pattern of plasticity can be maladaptive due to evolutionary mismatch.

The above approach analyzes individual decision making (both before and after HIREC) ignoring the game aspect (where expected payoffs depend on the actions of others) that can be important for many interactive (social or ecological) situations. In the HIREC context, game-theoretic models differ from the non-game approach outlined above in a key, under-appreciated manner. Rather than specifying an underlying mechanism (e.g., cue-response system) by which behavior is chosen, game theory typically specifies the expected payoffs (dependent on the behavior of the focal actor and the actions of others). These payoffs are then used to predict the frequency of each type of behavior, assuming that each individual is using a strategy that maximises expected payoff. Although this approach allows the evolutionarily stable strategies (ESSs) to be calculated [9], we identify that in many cases it will not allow the immediate effect of an environmental change to be predicted, because the decision mechanism is not specified. In the HIREC context, for mixed evolutionarily stable strategies, it is critically important whether the population consists of, for instance: 1) an ESS proportion of individuals with fixed strategies, or 2) individuals that each play the previously adaptive ESS proportion of behaviors, or 3) individuals that use a cue-based, optimally plastic behavioral mechanism to attain the ESS. We show this in relation to the simple and well-known hawk-dove game [10].

In this game context, we find that strikingly counter-intuitive results emerge, but for sensible reasons. First, if all individuals in a population exhibit optimal plasticity (i.e., if they adjust their strategy to fit the new ESS after HIREC), they can have lower mean payoffs than populations that exhibit less plastic strategies that do not adjust to match the new ESS. In addition, in a game context, when individuals adjust optimally to HIREC, changes that *increase benefits* (i.e. resource value) can *decrease expected payoffs* to all individuals. Additionally, HIREC that increases contest costs can increase expected payoffs. Below, we show when these counter-intuitive phenomena do or do not occur, and explain why they occur.

## Analysis

### The Hawk-Dove Game

The hawk-dove game (also known in other contexts as Chicken, or the snowdrift game) assumes that when competing for a resource, some individuals play the ‘hawk’ strategy (of aggressively competing for the resource) whilst others play ‘dove’ (taking the resource if it is not contested; so, on average, splitting the resource with other doves). When a hawk meets a dove, the hawk gains the resource (assumed to be of value *V*); however, when a hawk meets another hawk, there is assumed to be a cost, C, paid during the ensuing competition. The expected payoffs are shown in Table 1.

**Table 1:** Expected payoffs (to the row-player) in the hawk-dove game.

|  | Hawk | Dove |
| --- | --- | --- |
| Hawk | $(V - C)/2$ | $V$ |
| Dove | $0$ | $V/2$ |

Assuming that C > V (e.g., if competition is likely to result in serious injury), then in a population that consisted only of hawks, it would be better to play dove. Similarly, in a population consisting only of doves, a hawk would do extremely well. Thus there is an optimal frequency at which the strategies should occur; this evolutionarily stable strategy (ESS) occurs where the two strategies have the same expected gain in reproductive value. Denoting the probability of an individual playing hawk by *h*, the expected payoff of playing hawk is: (1 – *h*)*V* + *h*(*V* – *C*)/2, and the expected payoff of playing dove is: (1 – *h*)*V*/2. Setting these equal, we find the optimal probability of playing hawk, *h* = *V*/*C*, which is evolutionary stable strategy (ESS) when *V* < *C* [10].

### Consequences of mechanisms which determine phenotype post HIREC

It is well-known that the balance of frequencies between behaviors in the hawk-dove game could come about in numerous ways, even within a single species; e.g., via a genetic polymorphism where each individual is either a highly aggressive hawk or an unaggressive dove (i.e., with an ESS proportion being born as hawks), or through each individual being behaviorally variable [11]. However, the effect of these possibilities may be very different following HIREC. Assume for instance that before HIREC, the reward for an uncontested resource was *V* = 1, and the cost of a contest was C = 2. This means that half the individuals should behave as hawks (before the environmental change), with an expected payoff (to any individual) of 0.25 before HIREC (the calculations supporting this value and others in this text are detailed in Supplementary Information 1). We now consider the effect of HIREC decreasing the value of rewards, say from *V* = 1 to 0.5. This could happen, for example, if human disturbance reduces the quality of a contested territory or food resource in ways that are not transparent to the competitors. What would be the effect?

If the behavioral differences were the result of genetic polymorphisms then, immediately after HIREC (i.e., before the frequency of hawks and doves adjusts), the hawks would get a negative payoff following HIREC, of -0.125, whereas the doves would still get a positive payoff, of 0.125. Consequently, only the doves would, on average, gain reproductive value in interactions.

If, instead, the same balance (of *h* = 0.5) had ancestrally come about by each individual randomizing their choice in each encounter then, following HIREC, each individual would get the same mean payoff of zero.

A third possibility is that individuals may have evolved in circumstances where their likelihood of acting as hawk had evolved in relation to the strength of food (or habitat) cue that they received (i.e., letting h for each encounter depend on the value *V*). Under these circumstances, the individuals would display a reaction norm to the food rewards, increasing the probability of playing hawk as the perceived reward increased. Under such circumstances, the optimal reaction norm is to play hawk with probability *h* = *V/C*, as described in the previous section; this would still achieve the same expected payoff pre-HIREC (of 0.25) but, immediately after HIREC, they would adopt the best possible behavior of now playing hawk on only 25% of occasions, resulting in a positive expected payoff of 0.1875. Although the rate of reproductive success would have decreased due to HIREC, each member of the population would still tend to accrue reproductive value with numerous interactions – and would do even better than the doves in the genetic polymorphism case (discussed above, where hawks did very badly after HIREC).

Figure 1 shows the general effect of modifying *V* (from 1) whilst *C* remains fixed (at 2), for each of the three mechanisms of choice. Although the reaction norm is the optimal response to the conditions by each individual in the population, and does best if the environmental change decreases *V*, if the value of resources were to increase (*V* > 1), the reaction norm would result in a lower expected payoff than under the other strategies. This is because the proportion of individuals playing hawk would have increased (linearly with the change in *V*) immediately after the change, resulting in many more costs being paid through conflict than under the other strategies.

**Figure 1:**
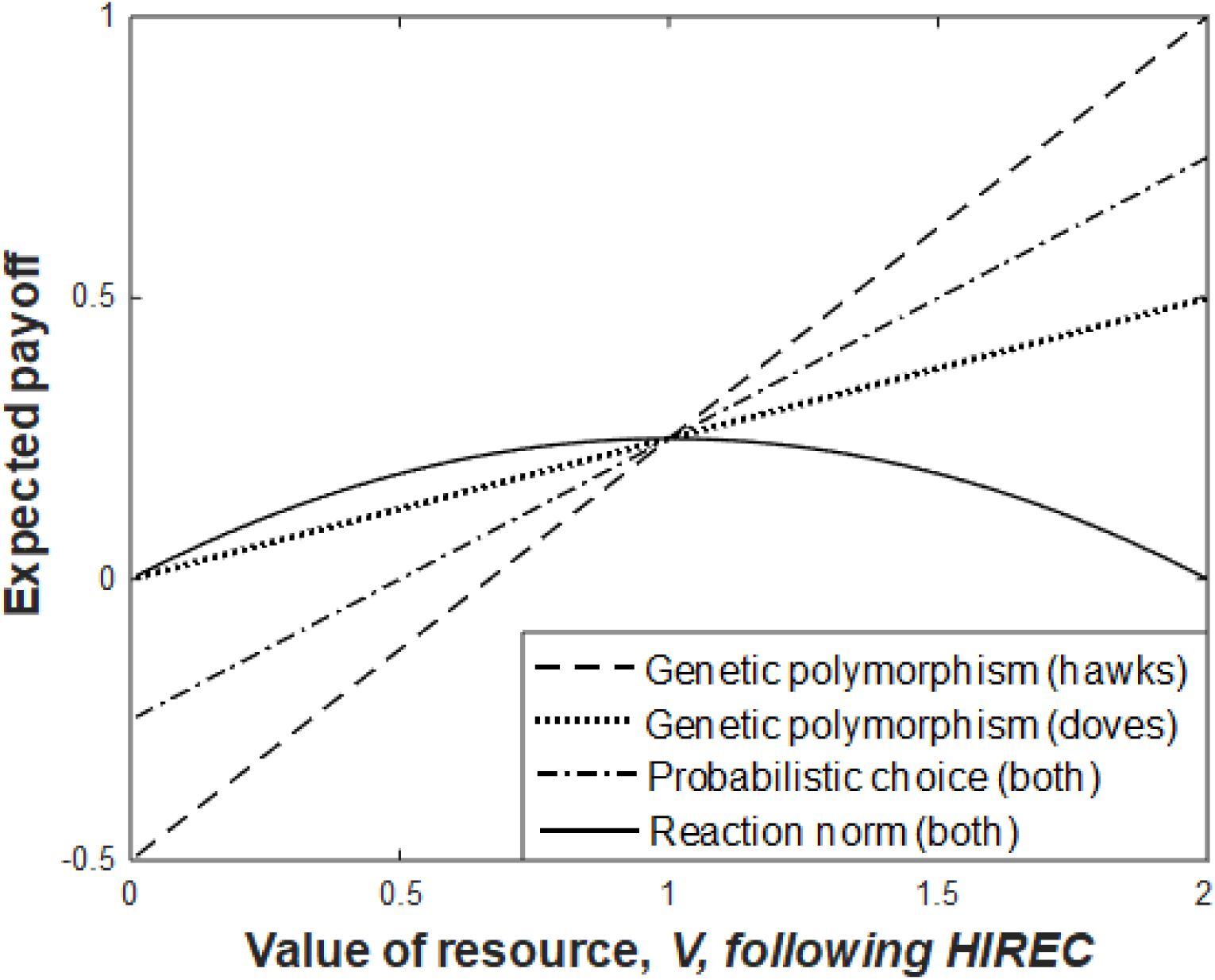
The evolved mechanism (supporting the pre-HIREC ESS) can have significant effects on payoffs immediately following HIREC. This shows the effect of altering *V* (from an initial value of 1) for genetic polymorphisms, probabilistic choice or an optimal reaction norm. Under the genetic polymoprhism and probabilistic choice mechanisms, the proportion of individuals playing hawk is assumed to remain the same (at 0.5) immediately after HIREC, whereas the reaction norm assumes that individuals reassess the situation in light of recognising the new value of *V* following the change, modifying their probability of playing hawk as *h* = *V/C*. Thus, under the reaction norm, the proportion of individuals playing hawk shifts linearly from 0 to 1 over the range of *V* shown. Note that this curve is invariant with initial *V*. Thus, if a population had an ancestral value of *V* greater than 1 (1.5 for instance) then, counterintuitively, small *increases* in the value of resources, *V*, would *decrease* expected payoff, and vice versa. [C = 2 throughout.]

**Figure 2:**
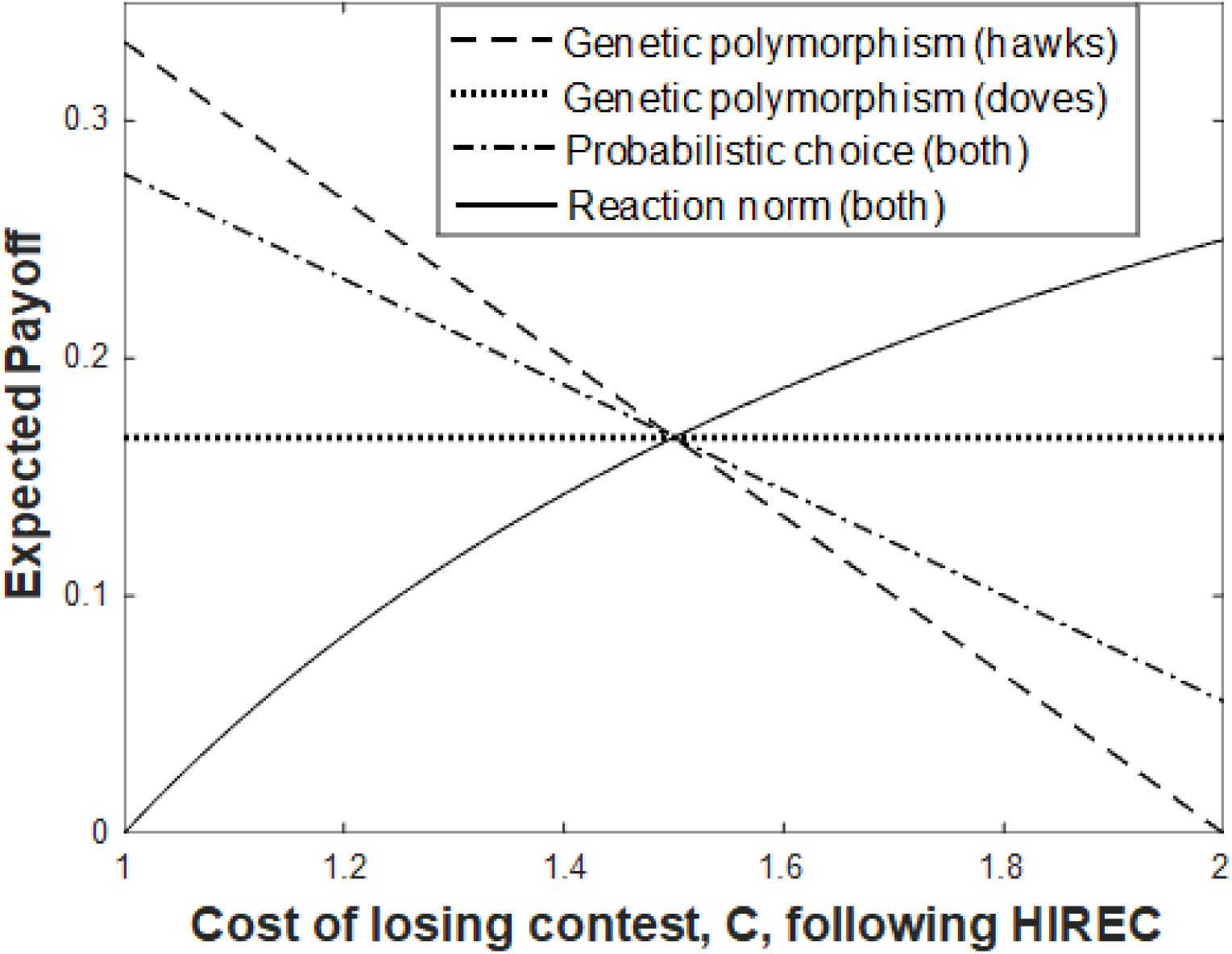
The effect of evolved mechanism (supporting the pre-HIREC ESS) on payoffs immediately following a change in the cost of contest. This shows the effect of altering *C* for genetic polymorphisms, probabilistic choice or an optimal reaction norm. Under the reaction norm, the expected payoff increases throughout the range as *C* increases – an opposite trend to that under the other strategies. Note that this reaction norm curve is invariant with initial *C*. [V = 1 throughout; initial (pre-HIREC) *C* = 1.5.]

Consider the situation when organisms that evolved plastic responses to changes in *V;* (i.e., so they shift their behavior adaptively in response to HIREC). If *V/C* starts below 0.5 then small changes in *V* have intuitive effects on fitness, but if *V/C* was greater than 0.5 before HIREC, then altering V will have a counter-intuitive effect: increasing resource value will lead to a reduction in expected payoffs and vice versa. The reason is that increased resource value results in a higher tendency to play hawk, thus more fights and ultimately a reduction in mean fitness. Conversely, reduced resource value reduces the tendency to play hawk, thus fewer fights, and an increase in mean fitness. In Supplementary Information 1 we also show that if HIREC increases the cost of conflict, *C*, then the expected payoff increases under the ESS (reaction norm) strategy; this occurs for any value of V between 0 and *C*.

In many intra-specific contexts, the mechanism must be specified (sometimes at several additional layers below that of the game) to carry out the analysis. This requirement to specify the mechanism of behavioral choice (as well as the functional payoffs), that we have shown in relation to the hawk-dove game, applies across a great many games. In Supplementary Information 2, we show that the mechanism would also need to be specified to consider the effects of HIREC in a producer-scrounger game [12]. The mechanism supporting the old ESS behavior (e.g., whether traits are fixed, probabilistically selected or dependent on cues) will affect subsequent population dynamics and timescales to reach a new equilibrium. We also identify in Supplementary Information 2 that a mechanism of decision would be required to predict outcomes following HIREC in an iterated Prisoner’s Dilemma game; thus the result does not only apply to games in which strategies have negative frequency dependent payoffs.

## Discussion

We have shown that altering the value of resources can have counter-intuitive effects. When individuals use a flexible rule that conforms to the ESS for the current conditions (i.e., the reaction norm, that shifts with conditions), then a decrease in resource value (e.g., a reduction in food magnitude) can increase expected payoffs, and vice versa. This effect occurs when the value of resources was previously at least half the expected cost of competition. Furthermore, increasing the cost of conflict in the hawk-dove game always increases the expected payoffs to individuals under the ESS strategy. In each case, the counter-intuitive results occur in populations using a reaction norm that follows the ESS, and the fact that this decrease in fitness occurs with increased resources (or decreased costs) can seem particularly counter-intuitive in the light that it occurs for individuals that would not be able to improve their strategy, even if they were able to choose a different strategy. This occurs because we are assuming that all individuals are using the same strategy; just like the tragedy of the commons [13], if only one individual were to adapt their behavior to the current conditions then that individual would do better than the others. But because all individuals are using the same strategy, an increased direct benefit (of the resource value increasing) can have indirect costs (through more individuals choosing the hawk behavior) that outweigh the direct benefits.

Why does the expected payoff increase and then decrease with as the value of the resource, *V*, increases? In cases where the resource value is low, relatively few individuals are behaving as hawks, and increasing *V* increases the direct benefits more than the indirect costs. This is because the probability of meeting a hawk increases linearly with *V* (for *V* between 0 and the cost of a contest, *C*). However, when *V* > *C*/2, more than half the population are already hawks, so an increase in *V* means that the corresponding linear increase in the number of hawks imposes more indirect costs than the direct benefits of the increase. In the case of increasing *C*, the indirect benefits always outweigh the direct costs.

Counter-intuitive effects in games are not uncommon—which is one of the reasons that models of game situations are so useful. For instance, Braess’s paradox [14] and the well-known Prisoner’s Dilemma provide examples where the Nash equilibrium that is reached by individuals does not achieve the maximum possible payoff to individuals. Nevertheless, the ease with which these effects occur in a situation as simple as the hawk-dove game suggests that they may not be uncommon in real situations. Furthermore, the different proximate details (about how the behavioral ESS is supported) show that the details of mechanisms can even result in differences over whether payoffs will increase or decrease as the value of resources, or the cost of conflict, increase.

We have also shown that to calculate the immediate effects of a change in a game-theoretic situation, additional assumptions will typically need to be introduced (on top of the assumptions of the original model and those of the environmental change), to know the degree of adaptive plasticity in the evolved organism. This contrasts with recent work on HIREC scenarios that do not involve game theory, which have used existing optimality models to set behavioral parameters before identifying how the system would respond following an environmental change (e.g., [6, 15], with the general approach specified in [8]).

In both game-theoretic and non-game scenarios, the mechanism being used pre-HIREC is fundamental to what will occur following HIREC – but the mechanism is often not specified (or known) in game theoretic scenarios. Simply knowing the pre-HIREC evolutionarily stable balance does not allow us to identify outcomes following HIREC without additional assumptions (or inferences) about the proximate drivers which maintained the pre-HIREC behavioral choices (this is even without the complications of learning and feedbacks – for instance of hunger potentially driving more hawk-like behavior – which would seem bound to further complicate what would occur following HIREC). There is a simple answer about mechanism in some circumstances though: when different species are involved, it can often be assumed that the differences are species-specific. For instance, [16] puts forward a model for how different bee species may respond to HIREC, in relation to competition for nesting space, where larger individuals have a competitive advantage. Because this is done on the basis of the individuals being of different species (like having genetic hawks and doves, rather than individuals choosing which type to play), the mechanism of difference – and thus decision – is clear.

The effect applies in both negative- and positive-frequency dependent cases. There are other games which are not frequency-dependent in either way. The war of attrition [9] is one well-known case, where the ESS is a mixed strategy for how long to persist in competition. It is easy to see in such a game also, the effect of HIREC will depend on additional assumptions that relate not to how the organism has evolved to respond to environmental cues. If an organism has evolved a strategy that does not pay attention to the payoff size (e.g., in a world where payoffs are relatively static) then their immediate behavior following HIREC will not be affected by the reward sizes having altered. If, instead, their strategy is dependent on the reward magnitude, then their response will immediately shift with the environmental change. It is also worth noting that in the war-of-attrition, under the ESS strategy (following the optimal reaction norm to food value), HIREC would result in no change to expected payoff, of zero. The war-of-attrition may therefore be one of the boundary cases between games where expected payoff decreases as the value of resources increase and those where payoff increases as resources increase (see [17]).

We have analysed cases as though the mechanisms of choice are thoroughly distinct but behaviors will typically be partially genetic, partially learned/ontogenetic, and partially cue-based [18]. The long-term outcomes in each case are similar (still governed by the functional payoffs of the game) if we assume that evolution will optimize the behavior (the ‘behavioral gambit’; [19]), but the behavioral adjustments and success of species following an environmental change can be quite different. The behavioral outcome immediately after an environmental change will depend on the evolutionary history of problems faced by the focal species and the range of responses they have evolved to cope with these problems. This raises an important question of the extent to which we can make reliable predictions about HIREC effects if we only know the balance between behavioral types, rather than the driving causes of those behaviors. Game theory is useful for understanding evolved (ESS) behaviors in an ancestral environment. We have shown that knowing only the probabilities of behaviors, without additional knowledge (or assumptions) of their causal mechanism, it is not possible to identify the effect of HIREC even in the initial interactions that immediately follow the environmental change. This even applies to whether environmental changes in simple scenarios (such as the hawk-dove game) will have positive or negative effects on populations. This in turn shows that the functional perspective that is taken in game theory is not sufficient to make predictions about the immediate effects of HIREC. Thus, to make worthwhile predictions about the effects of HIREC, researchers will also need to focus on the proximate pillars identified by Tinbergen [20], of mechanism and ontogeny.

## Acknowledgements

PCT and BJB were supported by the NSF (IOS 1456724 grant to AS). Thanks to John McNamara and Ken Binmore for useful comments.

## Author contributions

PCT had the formative idea and wrote the first draft of the ms. BJB, RM and AS contributed helpful suggestions along the way and improved the ms.

## Data availability

This study uses no data. Thus all data are readily available (or never available, depending on perspective).

## References

1. Sih, A., Ferrari, M.C.O., Harris, D.J. Evolution and behavioural responses to human-induced rapid environmental change. Evolutionary Applications 4, 367–387 (2011).

2. Candolin, U. & Wong, B. Behavioural Responses to a Changing World: Mechanisms and Consequences. (Oxford University Press 2012)

3. Tuomainen, U. & Candolin, U. Behavioural responses to human-induced environmental change. Biol Rev 86, 640–657 (2010).

4. Schlaepfer, M.A., Runge, M.C. & Sherman, P.W. Ecological and evolutionary traps. TREE 17, 474–480 (2002).

5. Robertson, B. A., Rehage, J. S. & Sih, A. Ecological novelty and the emergence of evolutionary traps. TREE 28, 552–560 (2013).

6. Trimmer, P.C., Ehlman, S.M., Sih, A. Predicting behavioural responses to novel organisms: state-dependent detection theory. Proc R Soc B 284, 20162108 (2017).

7. Sih, A. Understanding variation in behavioural responses to human-induced rapid environmental change: a conceptual overview. Animal Behaviour 85, 1077–1088 (2013).

8. Sih, A., Trimmer, P.C., Ehlman, S.M. A conceptual framework for understanding behavioural responses to HIREC. Current Opinion in Behavioral Sciences 12, 109–114 (2016).

9. Maynard Smith, J. Evolution and the Theory of Games. (Cambridge University Press 1982) 10

10. Maynard Smith, J. & Parker, G.A. The logic of asymmetric contests. Anita. Behav. 24, 159– 175 (1976).

11. Maynard-Smith, J. & Price, G. R. The logic of animal conflict. Nature 246(5427), 15–18 (1973).

12. Barnard, S.J. & Sibly, R.M. Producers and scroungers: a general model and its application to captive flocks of house sparrows. Animal Behavior 29(2), 543–550 (1981).

13. Lloyd, W.F. Two lectures on the checks to population. (Oxford University 1833).

14. Steinberg, R. & Zangwill, W.I. The prevalence of Braess’ paradox. Transportation Science 17(3), 301 (1983).

15. Crowley, P.H., Trimmer, P.C., Spiegel, O., Cuello, W.S., Sih, A. Predicting habitat choice after rapid environmental change. (Submitted)

16. Higginson, A.D. Conflict over non-partitioned resources may explain between-species differences in declines: the anthropogenic competition hypothesis. Behavioral Ecology and Sociobiology 71, 99 (2017).

17. Kaznatcheev, A. Online manuscript: https://egtheory.wordpress.com/2012/03/14/uv-space/ (2012).

18. McNamara, J.M., Dall, S.R.X., Hammerstein, P., Leimar, O. Detection vs. selection: integration of genetic, epigenetic and environmental cues in fluctuating environments. Ecol Lett 19, 1267–1276 (2016).

19. Fawcett, T.W., Hamblin, S., Giraldeau, L.-A. Exposing the behavioral gambit: the evolution of learning and decision rules, Behavioral Ecology 24, 2–11 (2013).

20. Tinbergen, N. On aims and methods in ethology. Zeitschrift fur Tierpsychologie 20, 410–433 (1963).

